# The Harsh Microenvironment in Early Breast Cancer Selects for a Warburg Phenotype

**DOI:** 10.1101/2020.04.07.029975

**Authors:** Mehdi Damaghi, Jeffrey West, Mark Robertson-Tessi, Liping Xu, Meghan C. Ferrall-Fairbanks, Paul A. Stewart, Erez Persi, Brooke L. Fridley, Philipp M. Altrock, Robert A. Gatenby, Peter A. Sims, Alexander R. A. Anderson, Robert J. Gillies

## Abstract

The harsh microenvironment of ductal carcinoma *in situ* (DCIS) exerts strong evolutionary selection pressures on cancer cells. We hypothesize that the poor metabolic conditions near the ductal center foment the emergence of a Warburg Effect (WE) phenotype, wherein cells rapidly ferment glucose to lactic acid, even in normoxia. To test this hypothesis, we subjected pre-malignant breast cancer cells to different microenvironmental selection pressures using combinations of hypoxia, acidosis, low glucose, and starvation for many months, and isolated single clones for metabolic and transcriptomic profiling. The two harshest conditions selected for constitutively expressed WE phenotypes. RNA-seq analysis of WE clones identified the transcription factors NFкB and KLF4 as potential inducers of the WE phenotype. NFкB was highly phosphorylated in the glycolytic clones. In stained DCIS samples, KLF4 expression was enriched in the area with the harshest microenvironmental conditions. We simulated *in vivo* DCIS phenotypic evolution using a mathematical model calibrated from the *in vitro* results. The WE phenotype emerged in the poor metabolic conditions near the necrotic core. We propose that harsh microenvironments within DCIS select for a Warburg phenotype through constitutive transcriptional reprogramming, thus conferring a survival advantage and facilitating further growth and invasion.

## Introduction

Ductal carcinomas *in situ* (DCIS) of the breast are a heterogeneous group of neoplastic lesions confined to the lumens of breast ducts. In early intraductal cancers, hyperplasia forces cells to grow towards the ductal lumens, which moves cells further from their supplying blood vessels that are restricted to the surrounding stroma (**Figure 1A**) ^1^. As a consequence, these cells are significantly nutrient-deprived. Hyperplastic tissue in DCIS can be > 0.16 mm thick, which is larger than the diffusion distance of oxygen in tissues, rendering the periluminal areas of DCIS hypoxic ^2,3^. This lack of oxygen would induce glucose fermentation due to a Pasteur effect, and the resulting production of lactic acid would make the periluminal areas of DCIS profoundly acidic. This has been verified following identification ^4^ and subsequent validation ^5^ of membrane-associated Lamp2b as a marker for acid-adaptation, which is abundant in the periluminal cells of DCIS. These microenvironmental properties of hypoxia, acidity, and nutrient deprivation exert strong selection pressure on cancer cell survival, and the metabolic adaptations subsequently feed back to the microenvironment, creating a dynamically changing landscape. Over many years in this environment, cells adapt and emerge with flexible, aggressive, and de-differentiated phenotypes^6^.

**Figure 1:**
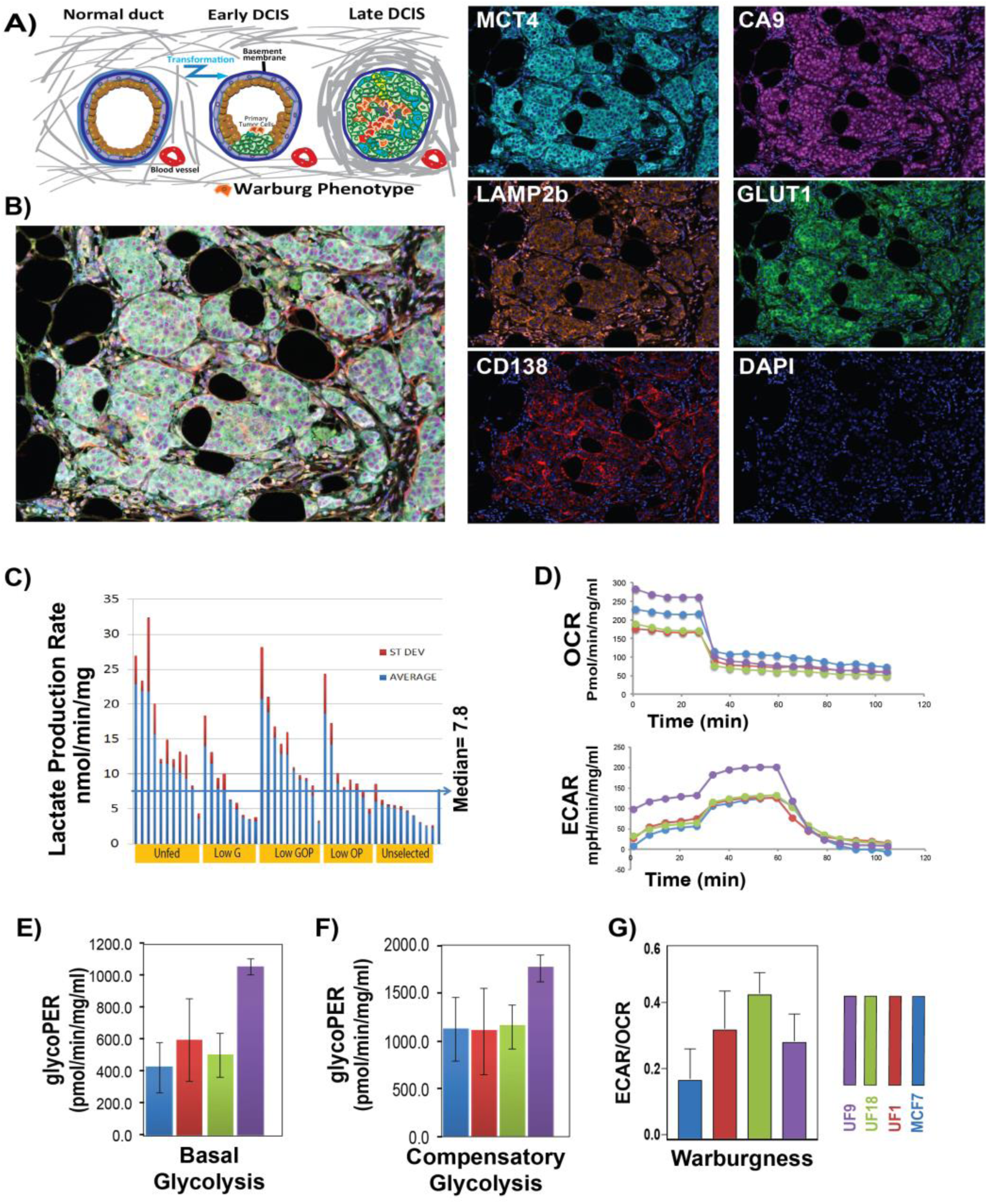
Early DCIS conditions can select for glycolytic phenotype. **A)** Schematic of early and late ductal carcinoma in situ (DCIS) progression. **B)** Multiplex IHC staining of DCIS patient sample with markers of glycolysis (Glut1[green]), acid adaptation (LAMP2b [orange]), hypoxia (CA9 [purple]), Lactate production (MCT4 [-cyan]), vasculature marked (CD138-red), and Nuclei (DAPI [blue]). **C)** Lactate production rate of clones grown out from cells selected under conditions of being selected through multiple rounds of the following conditions: unfed for 1 month (UF), low Glucose (G), low Glucose Oxygen and pH (GOP), low Oxygen and pH (OP), and growth in rich media (Control). **D)** Seahorse glycolytic rate assay to measure extracellular acidification rate (ECAR) and oxygen consumption rate (OCR) following addition of glucose. **E)** Basal Glycolysis was higher in UF cells but **F)** compensatory glycolysis showed no difference between control clones and overall UF clones. **G)** UF clones have higher WE phenotype (expressed as ECAR/OCR ratio) than control.

The most prominent metabolic hallmark to emerge from DCIS selection is the Warburg Effect (WE) phenotype, which is defined as aerobic glycolysis, where glucose is fermented to lactic acid even in the presence of adequate oxygen, contributing to the acidity of the ductal microenvironment ^1^. A WE is commonly observed in aggressive cancers ^7,8^ and has been exploited clinically with ^18^FDG-PET scans as a diagnostic marker of tumor stage and is prognostic of cancer outcome ^9^. Despite its almost ubiquitous expression in cancers, the causes and consequences of a WE remain a mystery. There have been dozens of mechanisms proposed, yet none have been proven. We have previously proposed that these conditions (hypoxia, acidosis, or nutrient deprivation) would select for cells with WE phenotype. In an initial study, cells were selected with periodic hypoxia (16 hr 0.2% O_2_, 8 hr 21% O_2_ for 50 cycles). Multiple clones were derived from surviving cells, and these were shown to be pan-therapy resistant, had an E-cadherin to N-cadherin switch, and a loss of p53, with a moderate increase in aerobic glycolysis that was not sustained^10^. In a subsequent study, we adapted cells to growth in acidic conditions, and this selected for a number of important phenotypes, including anchorage-independent growth, yet it did not select for cells with aerobic glycolysis, although cells adapted to acid pH did ferment glucose more rapidly at a low pH, compared to non-adapted cells^4,11,12^. Vogelstein’s group has shown that nutrient deprivation, specifically limiting (0.2 mM) glucose, promoted the outgrowth of pancreatic cancer cells that express mutant k-ras and a WE phenotype in mixed starting cultures, although *de novo* selection was not shown^13^.

Hence, we hypothesize that, if conditions in DCIS select for constitutive aerobic glycolysis, it may involve a complex and dynamic interplay between the multiple factors of hypoxia, acidosis, and nutrient deprivation. To test this, we subjected pre-malignant breast cancer cells to a series of these combined selection pressures, over a period of many months. At endpoints, individual clones were isolated and characterized for their metabolic and transcriptomic profiles. The resulting selected clones were enriched for populations that constitutively expressed an aerobic glycolytic (WE) phenotype. Transcriptomic analyses identified a number of relevant factors that could account for constitutive glycolysis, including SP1/KLF4 and NFкB. KLF4 expression was validated on selected clones using Western blots and immunocytochemistry (ICC). Tissue micro array (TMA) and whole mount staining of DCIS patients showed increased expression of KLF4 in DCIS samples, when compared to adjacent normal, as well as a relatively elevated expression at the core of each DCIS where the most selective environment exists. NFкB was also validated to be over-expressed at the protein level *in vitro*, with a significant increase in the transcriptionally active phosphorylated form, which was associated with increased HK2 expression. Knockdown of NFкB -related p65 reversed the WE in highly glycolytic clones.

We further investigated the emergence of Warburg phenotypes, in areas under harsh selective pressures, by adapting a previously published mathematical model of tumor metabolism and growth, informed by empirical data^14,15^. The model simulates a tumor growing in a homeostatic tissue, initialized within a ductal structure with diffusion-limited nutrients. Different tumor phenotypes were allowed to evolve due to selection, and multiple simulations showed that the selection of a Warburg phenotype occurred in the harshest conditions near the peri-luminal necrotic core. The model was calibrated to the *in vitro* results presented herein, and simulations under different conditions suggested that different modes of selection can be in action, depending on cellular turnover and the specific microenvironmental conditions. In particular, the harsh conditions had bottleneck-like selection events, whereas the less harsh conditions tended to show phenotypic drift.

Thus, we conclude that the microenvironmental conditions existing in DCIS are sufficient, with time, to select for cells with a WE phenotype. In this particular case, the switch to a WE phenotype is related to KLF4 as a phenotypic switch and/or NFкB expression as a survival strategy. This study unravels the role of harsh microenvironmental selection pressures in driving activation of pathways, controlled by key transcription factors, that lead to the WE phenotype and subsequent cancer progression.

## Results

### Harsh microenvironments, similar to early DCIS conditions, select for clones with higher aerobic lactate production rate

In early carcinogenesis, intra-ductal hyperplasia leads to significant alterations in the physical microenvironment, especially in peri-luminal cells that are far (>0.16mm) from their blood supplies; leading to a highly selective microenvironment of hypoxia, acidosis, and severe nutrient deprivation^1,6,16^. This suggests that the periluminal cells should be oxygen-deprived, which is consistent with increased expression of hypoxia inducible factor (HIF) client proteins, such as CA9 and Glut1, in periluminal areas of late-stage DCIS ^17,18^. As proof of principle we performed multiplexed immunohistochemistry (mIHC) on our DCIS stage patient whole mount samples for markers of high glycolysis (Glut1), hypoxia-induced acid production (CA9), non-hypoxia-induced acid production (MCT4), and acid resistance (LAMP2b) (**Figure 1B**). Our results illustrate that all these conditions exist inside the DCIS ducts individually or in combination. To better understand the impact of these conditions in DCIS breast cancer and their correlation to the WE phenotype, we subjected non-malignant MCF-7 cells to a range of selection forces, such as acidity (pH 6.7), hypoxia (1% O_2_), low glucose (0.1 mM), and combinations thereof, reflecting increasing levels of stress: i.e. low glucose (G), low oxygen and pH (OP); and low glucose, oxygen, and pH (GOP). Additionally, as an extreme condition, we selected cells by placing them in a flask and not replenishing the media for four weeks (unfed) which caused >95% of the cells to die (**Figure 1C**). We excluded acidosis and hypoxia alone as selection pressures, because previous results showed that these conditions alone do not strongly select for a WE phenotype^4,10^. Each of these harsh microenvironments resulted in significant cell death, followed by re-growth under rich microenvironmental (neutral pH, 21% O_2_ and 5.8 mM glucose) conditions, this process was repeated multiple (2-6) times with flasks re-gaining confluence, typically within 4 weeks, before re-exposure to harsh conditions. After the final outgrowth, we isolated individual clones (>20 per condition) both from controls that were continuously grown in a rich microenvironment and those that were selected to survive in harsh microenvironments (G, OP, GOP, unfed) by seeding individual cells in 96 well plates, which were then re-grown under rich microenvironmental conditions. These clones were then expanded in individual T25 flasks, which were then harvested for freezing and for metabolic profiling for rates of lactate production and glucose consumption under normal culture conditions, as the first sign of WE phenotype, using colorimetric kits. **Figure 1C** shows the lactate production rates (LPR, in nmol/min/mg protein) for individual clones from the 4 harsh and 1 rich (control) conditions. These data demonstrate that harsh environmental conditions preferentially select for clones with increased rates of aerobic lactate production. - production; specifically, the unfed and low GOP conditions had the greatest number of high lactate production rate (LPR) clones (**Figure 1C**). To relate our finding of high lactate production rate to the WE, we further measured lactate production and glucose consumption rates at the same time using a multi-analyte system (YSI 2900, Yellow springs OH) in 96 well plates. For these studies, we used the three unfed clones (UF1, UF9, and UF18) with highest lactate production rates and three clones from the rich microenvironment MCF7 cells with low LPR. Results shown in **Figure S1** confirmed the higher LPR, observed by colorimetric assays, in the harsh compared to rich microenvironment clones.

To confirm the WE phenotype of the harsh microenvironmentally selected clones, compared to parental MCF7 cells, we performed the Seahorse glucose stress test (GST) assay to measure both basal and maximal glycolytic capacity of cells, as well as their respiratory capacity (See method for more details) (**Figure 1D**). We found all the unfed (UF) clones had higher basal glycolysis rate compared to control clones (**Figure 1E**), although their compensatory glycolysis was not different in general (**Figure 1F**). However, compared to their parental MCF7 clones all UF clones showed an increase in the ratio of extracellular acidification to oxygen consumption (ECAR/OCR), which is a measure of the WE phenotype (**Figure 1G**). These results indicated that the harsh microenvironmental conditions similar to those found in early DCIS select for a WE (aerobic glycolytic) phenotype, more specifically the combination of low glucose, low oxygen, and low pH or starvation provide the greatest selective pressure for a WE phenotype.

### Clonal evolution under DCIS microenvironmental conditions

These data have demonstrated that harsh microenvironmental conditions selected for cells with increased rates of aerobic glycolysis. Further, and slightly counter-intuitive, these selected cells maintained their WE phenotype even after being placed in abundant nutrients and oxygen conditions for multiple generations, i.e. exceeding 20 passages. To investigate the mechanisms leading to this stable (“hardwired”) phenotypic switch, we performed RNA sequencing (RNA-seq) analyses of our harsh and rich microenvironment selected single cell-derived clones (see Methods). Briefly, the harsh (unfed, GOP, OP, G) and rich clones were plated in 96 well plates and grown to confluence, from which RNA was extracted and sequenced using PLATE-seq ^19^(see methods). After filtering, 12,568 genes were used for further analysis. Unsupervised clustering of the RNA-seq data identified five distinct groups, that corresponded to each of the microenvironmental conditions (**Figure 2A, Figure S2A, and Figure S2B**).

**Figure 2:**
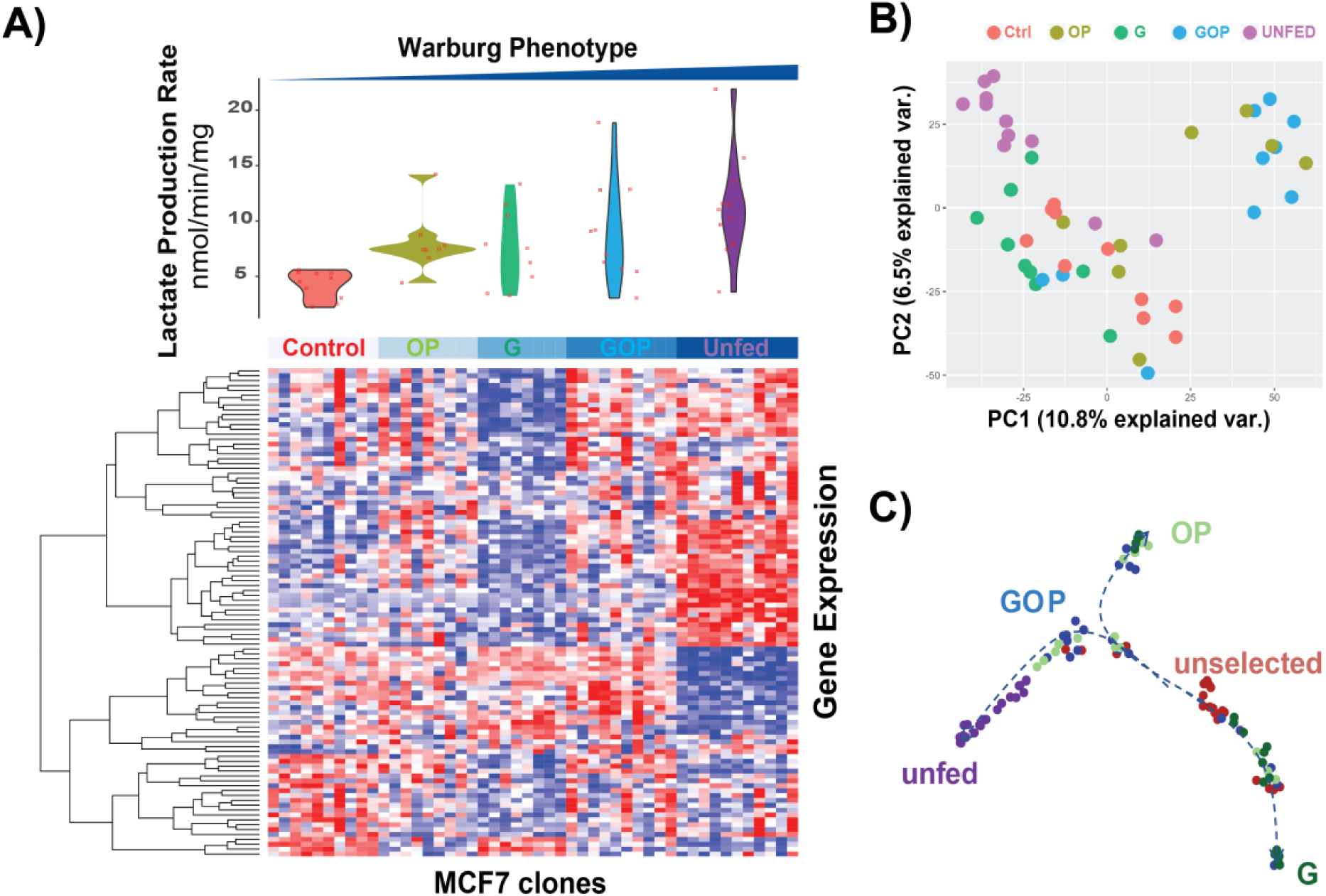
RNA sequencing analysis of selected clones reveals the molecular mechanism of switch to Warburg phenotype. **A)** Heatmap showing the top 500 most variable genes, grouped by selection condition. A preliminary analysis of RNA Sequencing data was performed by linearly regressing gene expression data with lactate production rate and filtered for significantly correlated or anti-correlated genes. Unsupervised clustering (**Figure S3**) of these data showed that 1000 most highly correlated and anti-correlated genes clustered within selection condition. **B)** Principal component analysis of gene expression data showed separation of the PE and GOP groups compared to control. **C**) Phenotype evolution trajectory alignment of single clone RNA-sequencing for evolving breast cancer cell populations. Cell fate analysis with Palantir was applied to the single clone RNA-sequencing dataset to determine differentiation potential from an initial, unselected, parental lineage to selected, phenotypic terminal states of G, OP, and unfed. UMAP projections were used to visualize the high-dimensional dataset and known identity of each clones was colored on the UMAP projections. Unselected clones were indicated in red, unfed clones were indicated in purple, G clones were indicated in green, OP clones were indicated in mint, and GOP clones were indicated in blue.

Principal component analysis (PCA) showed generally good segmentation for the different microenvironmental conditions (**Figure 2B and Figure S3)**. It is notable that the unfed (UF) cluster was readily segmented from the rest of the cells, suggesting that this condition, which more accurately reflects the *in vivo* situation, adds selection pressures beyond that imposed by the metabolic selections of G, OP, and GOP. Further, there was some overlap between the parental unselected and some of the selected (G, OP) clones, suggesting some clonal heterogeneity in the parental population or original phenotype recovery due to the highly plastic nature of MCF7 cells ^15^.

To determine which genes were associated with the WE phenotype, the gene expression data were linearly regressed against LPR using the *limma* and *voom* R packages ^20,21^. Six hundred seventy-six genes had a significant association with LPR (P_adj_ < 0.1), with 388 having a positive association (correlation coefficient > 0) and 288 having a negative association (correlation coefficient < 0) (**Figure 2A** and **Supplementary data1**).

To study phenotypic evolution under the different microenvironmental selection pressures on control MCF-7 cells, Palantir analysis (**Supplementary data 1**) was applied to the single clone RNA-seq dataset of unselected (parental), unfed, G, OP, and GOP clones to detect evolutionary trajectories of these clones and alignment along pseudo-time (**Figures 2C, S4, and S5**). All clones started in the rich microenvironment of the parental phenotype, indicated by Orange points in **Figure 2C**. Aligning clones along pseudo-time revealed three distinct terminal phenotypes: unfed, G, and OP. Interestingly, the GOP phenotype lay along the trajectory of the unfed terminus. Analysis of the differentiation potential of the clones showed that those aligned to the earliest pseudotime (dark blue in **Figure S4**) also had the highest differentiation potential, indicating that they are most likely to evolve to one of the terminal phenotypes over pseudotime. Likewise, especially for the unfed and G phenotypes, pseudotime was near 1 (yellow in **Figure S4**) indicating that these clones had the lowest differentiation potential and that they were at their terminal phenotypic states.

### Mathematical modeling shows the WE phenotype is rapidly selected in harsh conditions

To investigate the emergence of the WE phenotype in more detail, we extended a mathematical model of tumor metabolism^14,15^ to simulate the experiments presented herein. The extensions to the model that simulate the *in vitro* portions of this work are provided in the methods, Equation (1) and Tables 1 and 2. First, we applied our previously published model to simulate DCIS development *in vivo*. These results indicated that the WE phenotype primarily emerged from the most metabolically depleted area of a simulation, namely far from blood vessels and adjacent to the necrotic core. **Figure 3A-D** shows representative examples of these simulations (see **Figure S8** for more information about the simulated barcoding data in in **Figure 3B**). The WE phenotype is pink, and after 100 time increments (“days”) this phenotype began to emerge in the center of the duct where glucose and oxygen were highly depleted and the pH was acidic. This suggests that the harsh heterogeneous conditions of a tumor growing within a duct (or other similarly poorly vascularized region) would select for WE phenotypes.

**Table 1:** In vivo Parameters

| Param. | Value | Units | Description |
| --- | --- | --- | --- |
| $\delta x$ | 20 | $\mu m$ | Diameter of CA grid point |
| $p_D$ | 0.005 | 1/day | Normal tissue death rate |
| $p_\Delta$ | 0.7 | 1/day | Death prob. in poor conditions |
| $p_n$ | 5e-4 | 1/day | Necrotic turnover rate |
| $D_o$ | 1820 | $\mu m^2/s$ | Diffusion rate of oxygen |
| $D_g$ | 500 | $\mu m^2/s$ | Diffusion rate of glucose |
| $D_H$ | 1080 | $\mu m^2/s$ | Diffusion of protons |
| $O_o$ | 0.0556 | mmol/L | Oxygen concentration in blood |
| $G_o$ | 5 | mmol/L | Glucose concentration in blood |
| $pH_o$ | 7.4 | pH | pH of blood |
| $V_o$ | 0.012 | mmol/L/s | Max. oxygen consumption |
| $k_o$ | 0.005 | mmol/L | Half-max oxygen concentration |
| $k_G$ | 0.04 | mmol/L | Half-max glucose concentration |
| $k_H$ | 2.5e-4 | - | Proton buffering coefficient |
| $\Delta H$ | 0.003 | - | Pheno. variation rate (acid res.) |
| $\Delta G$ | 0.15 | - | Pheno. variation rate (glycolysis) |
| $A_d$ | 0.35 | - | ATP threshold for death |
| $A_q$ | 0.8 | - | ATP threshold for quiescence |
| $\beta_{T,min}$ | 6.1 | | Maximal acid resistance |
| $\beta_{G,max}$ | 50 | | Maximal glycolytic phenotype |
| $\beta_N$ | 6.65 | | Normal acid resistance |
| $\tau_{min}$ | 0.95 | Days | Min. cell cycle time |
| $\sigma_{min}$ | 80 | $\mu m$ | Min. vessel spacing |
| $\sigma_{mean}$ | 150 | $\mu m$ | Mean vessel spacing |
| $v_{mean}$ | 5 | - | Vessel stability |
| $p_{ang}$ | 0 | - | Angiogenesis rate |

**Table 2:** In vitro Parameters

| Param. | Value | Units | Description |
| --- | --- | --- | --- |
| S | 0.08 | - | Oxidative phosphorylation survival |
| $\Delta H$ | 0.005 | - | Pheno. variation rate (acid res.) |
| $\Delta G$ | 0.25 | - | Pheno. variation rate (glycolysis) |
| $A_d$ | 0.15 | - | ATP threshold for death |
| $A_q$ | 0.8 | - | ATP threshold for quiescence |
| $\beta_{T,min}$ | 5.8 | | Maximal acid resistance |
| $\beta_{G,max}$ | 10 | | Maximal glycolytic phenotype |
| $\beta_N$ | 6.2 | | Normal acid resistance |
| $\tau_{min}$ | 0.95 | Days | Min. cell cycle time |
| $k_G$ | 0.3 | mmol/L | Half-max glucose concentration |
| $k_{G0}$ | 2.5 | mmol/L | Baseline Half-max concentration |
| $k_o$ | 0.005 | mmol/L/s | Half-max oxygen concentration |
| $k_H$ | 1.2e-4 | - | Proton buffering coefficient |

**Figure 3.**
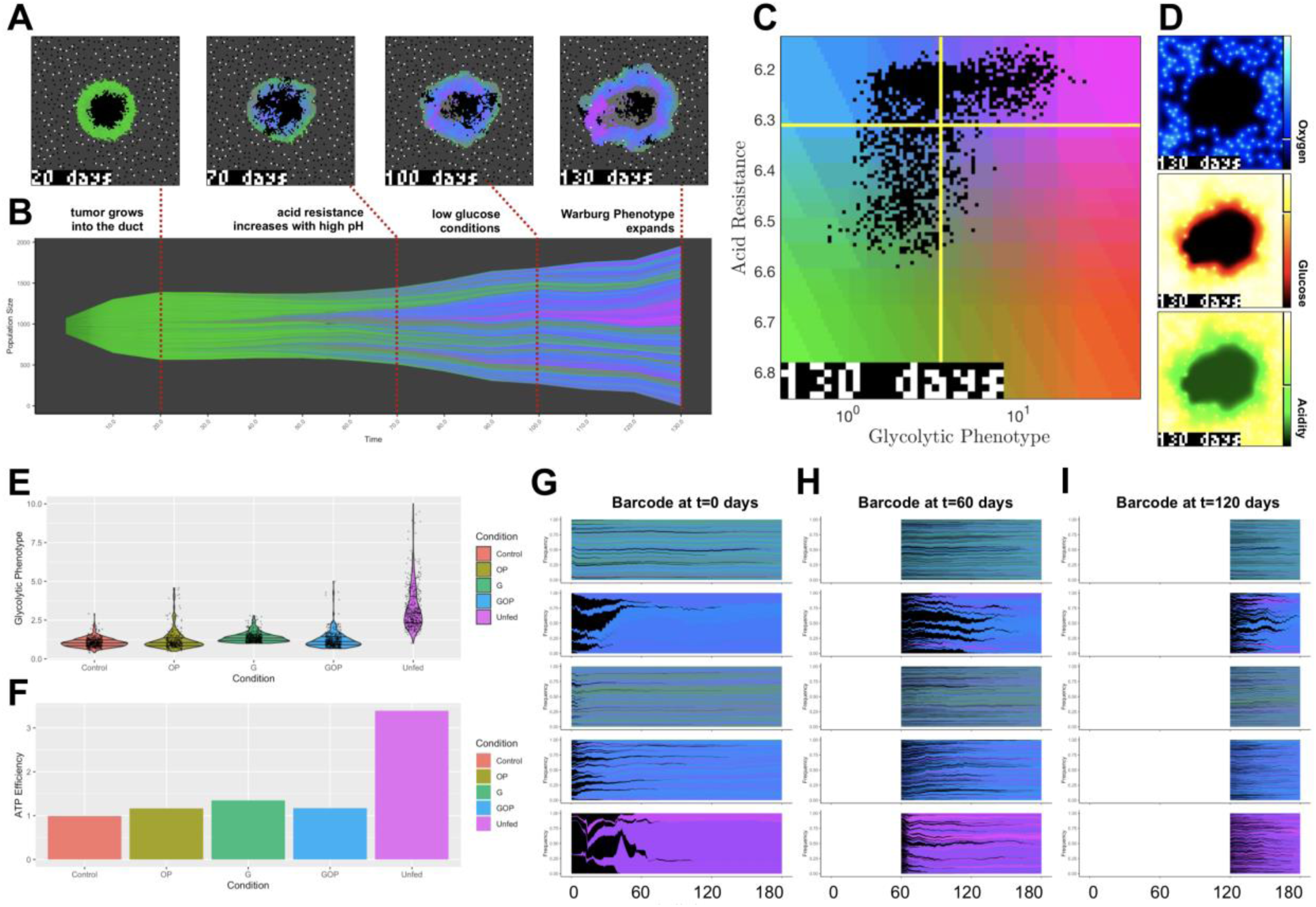
Computational modeling of the emergence of the Warburg phenotype. **(A-D)** A previously published 2-dimensional hybrid discrete-continuum homeostatic cancer metabolism model (see refs. 43, 44) shows the evolution of acid resistance (blue to green) and Warburg (blue to pink) phenotypes over time. The model simulates growth into ductal structure (A) where increased acidity in the center of the duct promotes acid resistance phenotypes (blue). After depletion of glucose, the Warburg phenotype emerges in harsh conditions near the center of the duct, on the edge of the necrotic core. (B) A Muller plot shows phenotypic selection and lineage over time. Vertical axis indicates size of clones, colored by its acid-resistance/Warburg phenotype shown in (C). Final distributions of oxygen, glucose, and acid are shown in (D). **(E-I)** An *in vitro* version of the model simulated for identical conditions as Figure 2A confirms that Warburg phenotype (E) emerges in harsh conditions (“unfed”). Furthermore, these cells have enhanced efficiency in producing ATP from nutrients (F). Model simulated barcode proportions are shown for 3 timepoints: 0 days (G), 60 days (H), and 120 days (I). Barcodes colored by average final phenotype with dead clones colored in black. Control and glucose-depleted conditions have low turnover, leading to slowed evolution. OP and Unfed conditions have increased turnover, selecting for Warburg phenotypes. Parameters (eqn. 1): S = 0.08, ko= 0.005, kg= 0.3, kg0= 2.5, V0= 0.93.

We then calibrated the model to directly simulate our *in vitro* experiments and the results of this simulation are shown in **Supplementary Video V1**. In short, to recapitulate an *in vitro* environment, blood vessels were removed from the simulation and nutrient concentrations and pH levels were changed globally across the ‘flask’. Cells were plated at the same seeding density as in the experiments and were allowed to adapt to their particular conditions through phenotypic drift, as in the original *in vivo* model. **Figure 3E-I** shows that the WE phenotype was selected primarily in the “unfed” case, when glucose and pH levels dropped significantly after 14 days without media change. The harsh conditions towards the end of the unfed period induced rapid turnover that enabled faster phenotypic drift. Furthermore, these cells exhibited increased ATP efficiency (**Figure 3E-F**), which is useful for survival in low glucose conditions.

The barcoding plots in **Figure 3G-I** show how phenotypic selection changes through the period of the simulation for each of the five conditions. The colors correspond to the phenotypes of **Figure 3C**, with pink cells having a WE phenotype. The unfed condition (bottom panel) showed rapid turnover of the population, driven by an early bottleneck, which quickly drove adaptation to the WE phenotype due to the severely depleted glucose and the highly acidic microenvironment. In contrast, the other conditions showed less turnover, even though some had low glucose and/or acidosis. The adaptation was slower and was primarily aimed at mitigating acidosis rather than becoming glycolytic. Notably, unfed cells with a WE phenotype emerged from only a few of the initial cells. This is in contrast to our metabolic profiling of the unselected clones, which showed that one clone had a slightly elevated lactate production rate, alluding to the likely pre-existence of this phenotype (**Figure 1C**).

### The role of transcription factors in selection of WE phenotype

To investigate whether specific transcription factors were involved in the transcriptional switch for generating a WE phenotype, we used the list of 388 significantly and positively associated genes for enrichment analysis by oPOSSUM and Enrichr^22,23^. oPOSSUM “single site analysis” with “human” was selected with default parameters. The top oPOSSUM hit was KLF4 (Z-score = 54.053) (**Figure 4A**). As a test of the oPOSSUM analysis, we investigated whether cells from UF clones had high nuclear KLF4 expression using ICC. **Figure 4B** shows nuclear localization and higher expression of KLF4 in UF clones compared to their parental MCF7 cells. KLF4 plays a role in early development and promotes a stem-like phenotype ^24-26^. Consistent with this, we also observed increased RNA expression of genes associated with stemness (Supplemental **Table S1**).

**Figure 4:**
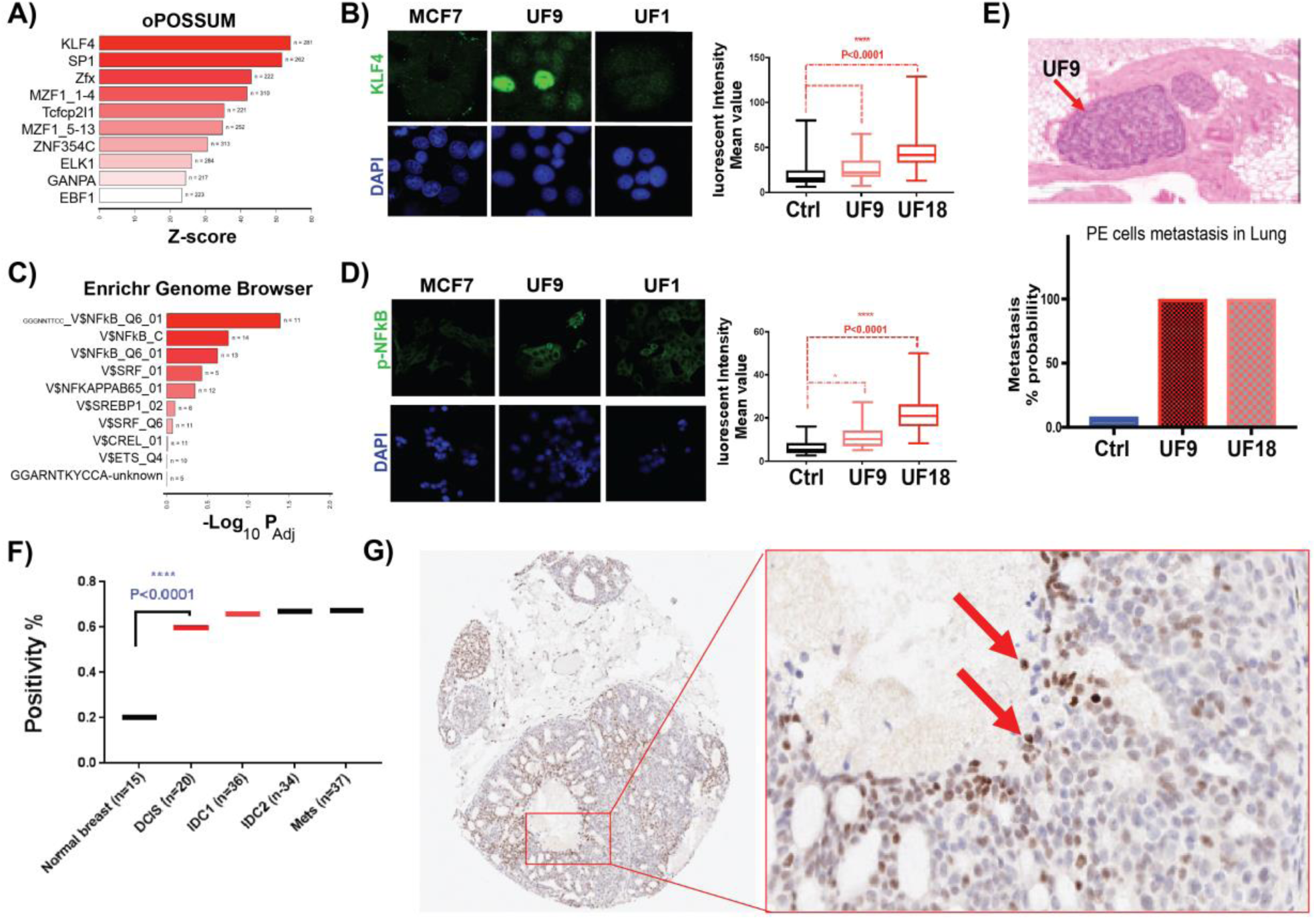
Clinical validation of KLF4 expression in breast cancer patients’ samples. **A)** Enrichment analysis of 388 genes positively associated with LPR using oPOSSUM revealed KLF4 as top regulator of lactate production rate (LPR) genes. **B)** Immunocytochemistry (ICC) analysis of UF and parental MCF7 validates the higher expression and nuclear localization of KLF4 in UF cells. **C)** Enrichment analysis NFкB as top hit in WE phenotype TF analysis. **D)** ICC validates the expression of p-NFкB in UF cells. **E)** Harsh condition in DCIS selects for aggressive cells that can invade other organs. 1 million of MCF7 parental, UF9 and UF clones were injected to tail vein of NSG mice and looked for possible metastasis. There was only one metastasis in the control group of twelve mice compared to 100% metastasis in both UF cells. **F)** TMA analysis of 204 breast biopsies of Moffitt cancer center patients for KLF4 expression shows higher expression of this protein in DCIS compared to adjacent normal tissues. The expression of KLF4 stays high with higher grade of breast cancer. **G)** Representative images of the TMA analysis done in A. The box is zoomed in the center of the duct to show spatial correlation of KLF4 in DCIS samples. Cells with high KLF4 expression are located to center of the duct in DCIS samples that proves our hypotheses that harsh condition selecting the cells with glycolytic phenotype through a transcriptional factor switch.

We also queried the Enrichr “Genome Browser PWMs” pathway, which contains a list of genes and associated binding motifs from transcription factors, and identified an NFкB-associated list as significantly enriched (P_adj_ = 0.04) (**Figure 4C**). To investigate this, we performed Western blotting on UF clones and parental MCF7. Supplemental **Figure S10A** shows that the total amount of NFкB was slightly, but significantly, higher in the UF clones, relative to control clones (C clones). In contrast, Western blot of phospho-NFкB (p-NFкB) shows that the p-NFкB, which promotes nuclear localization, was significantly elevated in UF clones, compared to that present in the C clones (Supplemental **Figure S10A**). To further investigate this, we performed ICC on UF clones and parental control MCF7 and found higher expression of p-NFкB in nuclei of UF cells (**Figure 4D**). To investigate the role of NFкB in promoting glycolysis in these systems, 3 separate anti-p65 siRNAs were prepared and tested for efficacy against UF-18 cells and were all able to effect knockdown at different doses. Using the p65siRNA-C, we were able to optimally knockdown expression in UF-18 and UF-9 clones (Supplemental **Figure S10B)**. In UF-18 cells, knockdown of p65 significantly reduced aerobic glycolysis (**Supplemental Figure S10C)** to the levels observed in non-selected cells. In addition to NFкB, other metabolically relevant proteins that were observed to be different between UF and C clones were pAkt and the NFкB client, HK2 **(Supplemental Figure S10A)**, which were also consistent with increased aerobic glycolysis in the UF cells. These results propose the pro-survival activity of NFкB in our unfed cells, which has also been shown for acid exposed cells in sarcoma ^27^.

To determine which of the cells expressing a WE were more aggressive compared to non-selected parental clones, we injected cells from two UF clones and 1 unselected clone into the tail vein of NSG mice and observed that 100% of both UF cell groups formed metastases, while only one mouse (8.3%) showed metastasis with MCF7 parental cells (**Figure 4E**).

#### KLF4 expression in clinical breast tumor samples

To further investigate KLF4 expression following selection of WE phenotype in DCIS lesions, we then interrogated expression of KLF4 in breast tumor samples from the Moffitt Cancer Center Tissue Micro Array (TMA) collection. KLF4 was selected for further clinical validation over NFкB based on its nature of switching phenotype in stem cells and early embryonic cells ^25^ as well as its previously shown role in regulating glycolysis in stem cells and pancreatic cancer ^28,29^. Breast TMAs from Moffitt breast cancer patients were used to study the expression level of KLF4 at different stages. TMA4 contains 204 biopsy cores including adjacent normal samples, DCIS, IDC with no metastasis, IDC with local metastasis, and lymph node metastasis core biopsy samples. Staining of TMA4 with KLF4 antibody showed significantly increased expression of KLF4 in DCIS samples compared to normal breast tissue. The KLF4 expression remained high in IDCs and metastasis samples (**Figure 4F and G, Supplemental figure S11**). To relate the role of KLF4 to selection of the WE phenotype in DCIS, we performed spatial analysis of KLF4 in our DCIS samples and observed that multiple sites had very high expression of KLF4 at the center of the duct, where access to nutrient resources is severely restricted, and thus exerting a strong selection pressure with regard to increased acidosis and decreased oxygen (**Figure 4G**).

## Discussion

The Warburg effect phenotype is associated with progression and aggressiveness of cancers and is defined by a high glycolytic rate in the absence or presence of oxygen (aerobic glycolysis). Most cancer cells reprogram their metabolism in favor of aerobic glycolysis despite the presence of plentiful oxygen in their microenvironment. This observation was first reported by Otto Warburg and is thus referred to as the “Warburg effect”^30-33^. We and others have observed this high glycolysis rate in tumors using positron emission tomography (PET)^34^. We also know that most cancer cells in hypoxic environments (Pasteur effect) compensate for the low ATP yield of glycolysis by overexpression of glucose transporters, such as Glut1^35^. The driving theory for why the Warburg effect takes place in cancer is that the high rate of glycolysis benefits cancer cells by increasing ATP production. It also provides many intermediates, that are used in subsidiary metabolic pathways for de novo synthesis of nucleotides, amino acids, lipids and NADPH, that are required for cancer cell survival and proliferation. However, none of these cellular regulations individually are enough to hardwire the Warburg phenotype in cells, because they can be altered based on microenvironmental conditions. At the center of individual DCIS the harsh microenvironment consists of low glucose, low oxygen, and a high acidity. Therefore, we hypothesized that there are some biological controls or switches at the genome, transcriptome, or epigenome level that initiate and control the Warburg phenotype.

Previous research has shown evidence of mutational drivers, such as p53 and KRAS, upregulating the Warburg phenotype in different cancer types, however, none of these mutation-driven Warburg phenotypes are consistently present in all cancer cells. Furthermore, there are cancer cells with a WE and no known driver gene mutations. This suggests that there may be mutation independent drivers of this phenomenon i.e. the microenvironment. To test our hypothesis, we probed the transcriptome of single selected clones under different microenvironmental conditions recapitulating the environments found in DCIS. Using single clones over single cells had the benefit of allowing us to measure the derived diversity and heterogeneity of a single cell’s progeny over time. Surprisingly, we found a highly variable transcriptome amongst the clones across all of the selection conditions, which may have been lost at the single-cell resolution. Using sophisticated transcriptome analysis of oPOSSUM and Enrichr, we discovered the transcription factor KLF4 controls all of the LPR genes. KLF4 was previously identified as one of the essential factors for iPS cell development^25^. KLF4 was previously reported to regulate WE phenotype^28,29,36^, although none of these studies connected the KLF4 expression or activation to microenvironmental conditions as evolutionary selection pressures. Here, we have shown that KLF4 induced WE is connected to the microenvironment of cancer cells in DCIS lesions. Open questions still remain regarding the heterogeneity of KLF4 expression in selected clones as well as clinical samples. This might imply redundant mechanisms, such as NFкB, that we also uncovered as a mechanism to maintain the WE phenotype or co-opt adjacent cancer cells^37^.

Finally, these results are paradoxical to our notion that cells under very low nutrient conditions should reduce their demand and energy expenditures based on the energy availability. Our findings suggest that the Warburg phenotype may be more efficient than previously assumed since we show that the WE phenotype is a highly regulated and controlled energy consumption source. Our results also illuminate the evolutionary trajectory of the Warburg phenotype driven by microenvironment selection pressures. We observed that transcription factors can activate the WE phenotype under appropriate environmental conditions that can both select for the WE phenotype and facilitate its hardwired statues. The activation of transcription factors, such as KLF4 and NFкB, may serve as an adaptive mechanism for cancer cells to switch to fitter phenotypes (Warburg phenotype) that can withstand the harsh environmental selective forces found in early DCIS lesions.

## Methods

### Cell culture and *in-vitro* clonal selections

MCF-7 cells were acquired from American Type Culture Collection (ATCC, Manassas, VA, 2007–2010) and were maintained in RPMI 1640 (Life Technologies, Cat# 11875—093) supplemented with 10% fetal bovine serum (HyClone Laboratories). Growth medium was further supplemented with 25?mmol?l^−1^ each of PIPES and HEPES and the pH adjusted to 7.4 or 6.7. Cells were tested for mycoplasma contamination and authenticated using short tandem repeat DNA typing according to ATCC’s guidelines.

### Western blotting

Selected and non-selected MCF-7 cells were grown with the same number of passages and used for whole-protein extraction. Lysates were collected RIPA buffer containing 1 × protease inhibitor cocktail (P8340; Sigma-Aldrich). Twenty micrograms of protein per sample was loaded on polyacrylamide–SDS gels, which later were electrophoretically transferred to nitrocellulose membranes. Membranes were incubated with primary antibodies against rabbit monoclonal KLF4 (1:1,000, ab215036 Abcam), NF-кB (1:1000, # 8242 Cell SignalingHK(1:1000,#2867s Cell Signaling), p-AKT (1:1000, #4060s Cell Signaling), Tubulin (1:1000, #3873 Cell Signaling) and GAPDH (1:4,000, antirabbit; Santa Cruz Biotechnology).

### siRNA Transfection

Three unique 27mer RELA human siRNA oligo dupliexes (SR304030A, B and C) were obtained from Origene (SR304030). Universal scrambled negative control siRNA duplex (SR30004, Origene) were used as a non-targeting control for this study. Cells were seeded in a six-well plate and reached 70% to 80% confluence before transfection. Cells were transfected with the negative control siRNA or p65-targeting siRNA according to standard protocols using lipofectamine RNAiMAX transfection reagent (13778030, Themo Fisher).

### Immunofluorescence

Selected and non-selected MCF-7 cells cultured with the same number of passages were rinsed with PBS, fixed in cold Methanol:Acetone (1:1) for 10 minutes and further permeabilized by 0.5% triton 100 and then blocked with 5% bovine serum albumin in PBS. Samples were incubated with KLF4 rabbit monoclonal primary antibody (1:100; ab 215036 Abcam) and secondary Alexa-Fluor 488 antirabbit (1:1000) antibody. Coverslips were mounted using ProLong Gold Antifade Reagent (Life Technologies) and images were captured with a Leica TCS SP5 (Leica) confocal microscope.

### Glycolytic and oxygen consumption rate measurements (Seahorse)

Glycolytic rate of MCF7 and selected MCF7 cancer cells was measured using Seahorse XF96 extracellular flux analyzer and a glycolysis rate kit (Seahorse Biosciences). Oxygen consumption rate (OCR) and extracellular acidification rate (ECAR) of cancer cells were determined by seeding them on XF96 microplates in their growth medium until they reached over 90% confluence. In these studies, seeding started with 20,000 cells (80% of well area). Measurements were determined 24 hour later when the cells reached the 90% confluence. One hour before the seahorse measurements culture media were removed and cells were washed 3 times with PBS followed by addition of base medium (non-buffered DMEM supplemented with 25 mM glucose) or our non-buffered only glucose containing solution. For glycolytic rate measurements, mitochondria inhibitors including rotenone (1μM) and antimycin A (1μM), were injected after basal measurements of ECAR and OCR of the cells under treatment to stop the mitochondrial acidification. 2-deoxy-glucose (100 mM) was added next to bring down glycolysis to basal levels. Finally, data were normalized for total protein content of each well using the Bradford protein assay (Thermofisher). Seahorse measurements were performed with 4-6 technical replicates and these experiments were repeated four times.

#### Solutions for seahorse experiments

2mM HEPES, 2mM MES, 5.3 mM KCl, 5.6 mM NaPhosphate, 11 mM glucose, 133 mM NaCl, 0.4 mM MgCl_2_, 0.42 mM CaCl_2_, titrated to given pH with NaOH. For reduced Cl-experiments, 133 mM NaCl was replaced with 133 NaGluconate and MgCl_2_ and CaCl_2_ were raised to 0.74 and 1.46 mM, respectively, to account for gluconate-divalent binding. Amount of dilute HCl or NaOH added to medium to reduce pH to target level was determined empirically.

Respiratory capacity is a measure of the maximum rate of O_2_ consumption and mitochondrial electron transport in a cell^38^. Glycolytic capacity is the maximum rate of glucose conversion to pyruvate and/or lactate by a cell. Glucose breakdown to two lactates produces two protons, allowing for the capability of indirect measurement of glycolytic rate using the extracellular acidification ^38^. Compensatory glycolysis is the maximum possible rate of glycolysis in cells following inhibition of oxidative phosphorylation with rotenone/antimycin. The WE phenotype (“Warburgness”) can be expressed as the ratio of glycolysis (ECAR) to oxidative phosphorylation (expressed as the oxygen consumption rate, OCR) from the GST.

### RNA-seq

High-multiplexed library preparation for RNA-seq (PLATE-seq) was performed as described previously ^19^. Briefly, we captured poly-adenylated mRNA from cell lysates using a 96-well plate with oligo(dT) grafted to the inner walls of each well (Qiagen). Next, we eluted the poly-adenylated mRNA and reverse transcribed using 96 different barcoded oligo(dT) primers (Integrated DNA Technologies). Following exonuclease digestion of excess primers, the barcoded cDNA libraries were pooled for second-strand synthesis and Illumina library construction. We sequenced the resulting pooled, barcoded, 3’-end libraries on an Illumina NextSeq 500.

### Metabolic Profiling

Cells were seeded in a 24-well plate in the growth medium containing 10% FBS under standard culture condition. Once cells reach 90% confluence, the growth media were removed and cells were washed twice in PBS and incubated in 2% serum and phenol-red free medium for 24 h. Then, the media were collected for lactate production assay. L (+)-Lactate was measured with a colorimetric kit (BioVision, K627-100) according to the manufacturer’s instructions. Absorbance (OD 450 nm) for each sample was background corrected with the culture medium (2% FBS) collected from the well without growing cells. Final data of the lactate production rates were normalized to the protein amount per well.

Lactate and Glucose concentration measurement was also done by YSI machine followed their protocol (YSI 2900 multi-analyte system (YSI, Yellow springs OH)).

### RNA Sequencing data analyses

#### Bioinformatics Processing and Statistical Analysis of RNA-Seq Data

Paired-end PLATE-seq data have the sample-identifying barcode sequences in read 1 and transcript sequences in read 2. We first aligned read 2 to the hg19 human genome with UCSC known genes transcriptome annotation using STAR39. Next, we demultiplexed the aligned reads based on the barcode sequence in read 1.Finally, we quantified the number ofuniquely alignedreads associated with each gene in each sample using the featureCounts ^39^.

We next filtered genes and only kept those with > 2 counts per million in at least 5 samples resulting in 12,568 genes for further analysis. To account for differences in library size between the samples, trimmed mean of M values (TMM) normalization was applied, followed by data transformation using the mean-variance relationship estimated on the observed log count data as implemented in the R package *voom*^20,40^. This results in approximately normally distributed count data for each gene, thus allowing for standard normal theory methods to be applied. We determined there that no batch effects were present using principal component analyses.

Association of gene expression with continuous measure of lactate production rates (LPR) was completed with linear regression models using the *limma* package ^21^. Genes with false discovery rate (FDR) q-values < 0.10 were considered significant^41,42^. Gene list enrichment was performed using oPOSSUM (http://opossum.cisreg.ca/oPOSSUM3/) and Enrichr (https://amp.pharm.mssm.edu/Enrichr/) ^22,23^.

### Evolutionary Trajectory Analysis

Alignment of cells along their evolutionary trajectories from the parental, unselected lineage to several selected stated was performed using Palantir ^43^, a recently published trajectory-detection algorithm for single cell RNA-seq data. Here, with single clone RNA-seq, we had complete RNA-seq expression per clone and did not need to impute any missing data, as is done with single cell RNA-sequencing datasets. Palantir models cell fate choices as a continuous probabilistic process over pseudotime, estimating the probability of a cell in an intermediate state to reach a terminal state (here: G, OP, and GOP). Palantir calculates differentiation potential of a given cell leveraging the entropy over branch probabilities. Differentiation potentials near 1 correlate with earlier in the pseudotime lineage, which in this case corresponds to the parental lineage (unselected), and indicate cells with the highest potential of become a different phenotype over time. The high-resolution data allows for mapping of gene expression trends and dynamics over this pseudotime, which can be interpreted as how gene expression changes as the populations were driven from the parental lineage (unselected) to alternate terminal trajectories of G and OP phenotypes. Visualization of this dataset was performed using UMAP projections ^44,45^ of the high-dimensional dataset and further analyses were overlaid onto this representation. Python code implementing Palantir on this single clone dataset is available in the Supplement.

### Mathematical Modeling

We used the mathematical model described in ^14^ and extended in ^15^ as a starting point. An interaction network and decision tree for the model are shown in **Figure S7**. For the *in vivo* simulations in **Figure 3**, panels A-D, we set up an initial condition of a duct, as in ^15^. Vascular was initialized with normal vascular density outside the duct and no vessels within. For the *in vitro* simulations in **Figure 3**, panels E-I, we altered the model as follows: vasculature was removed, and concentrations of oxygen, glucose, and protons (pH) were considered to be well-mixed, and therefore had a global value across the simulation domain. No diffusion was necessary, and metabolic reaction rates for glucose were calculated per cell and then summed across the entire population for each time step. This lowered the concentration of glucose over time; the pH was calculated via the metabolic equations of the model.

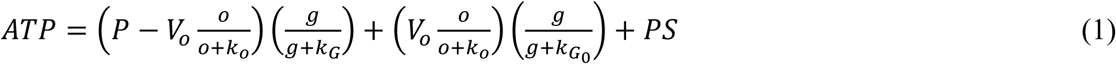

where P is the cell’s glycolytic phenotype while g and o are glucose and oxygen concentrations, respectively. Cells in the model were shown to survive in the unfed conditions well after glucose was depleted, suggesting that a secondary survival effect was in operation. This could be due to glutamine in the media, autophagic response, etc. To account for this behavior in the model, we added a term to the equation for ATP production under the hypothesis that this behavior emerges in concert with the Warburg phenotype. The term is a simple linear scaling with the glycolytic phenotype (*pG*) of a given cell, *kS pG*, added to the ATP production derived from normal metabolism (**Figure S9**). We fit the parameter *kS* based on the dynamics of the population and metabolites seen in the experimental system.

Replating was mimicked in the simulation by restoring the nutrient and pH values to their initial conditions every 3 or 14 days, as per the experimental protocol for the 5 different conditions. Simulations were implemented using the “Hybrid Automata Library” framework^46^ and barcoding visualized using the EvoFreq package in R^47^. Parameters for in vivo and in vitro models are below ^14,15^.

### Statistical Analysis -1

#### Bioinformatics Processing and Statistical Analysis of RNA-Seq Data

Primary analysis and de-multiplexing are performed using Illumina’s CASAVA software, resulting in de-multiplexed FASTQ files for subsequent analysis by the mapping software and aligner. These data will then be checked with *fastqc* program for quality assessment. Then *cutadapt* will be used to trim off adaptor contaminant sequences and low-quality bases at the ends. Reads pairs with either end too short (<25bps) will be discarded from further analysis. *Fastqc* will be used again to examine characteristics of the sequencing libraries after trimming and to verify its efficiency. Next, trimmed and filtered reads will be aligned to the hg19 human transcriptome using *STAR*^48^, followed by gene abundance estimation completed using *RSEM*^49^, as this approach accounts for reads mapping to multiple locations.

We next filtered genes and only kept those with > 2 counts per million in at least 5 samples resulting in 12,568 genes for further analysis. To account for differences in library size between the samples, trimmed mean of M values (TMM) normalization was applied, followed by data transformation using the mean-variance relationship estimated on the observed log count data as implemented in the R package *voom*^20,40^. This results in approximately normally distributed count data for each gene, thus allowing for standard normal theory methods to be applied. We determined there that no batch effects were present using principal component analyses. Association of gene expression with continuous measure of Lactate production rates (LPR) was completed with linear regression models using the *limma* package ^21^. Genes with false discovery rate (FDR) q-values < 0.10 were considered differentially expressed ^41,42^. Gene list enrichment was performed using oPOSSUM (http://opossum.cisreg.ca/oPOSSUM3/) and Enrichr (https://amp.pharm.mssm.edu/Enrichr/)^22,23^.

## Acknowledgements

We gratefully acknowledge funding from both the Physical Sciences Oncology Network at the National Cancer Institute through grant (U54CA193489) and the Cancer Systems Biology Consortium grant (U01CA232382) as well as support from the Moffitt Center of Excellence for Evolutionary Therapy. This work was also supported partly from R01 grant (R01CA077575). This work has been also supported in part by the Biostatistics and Bioinformatics Shared Resource at the H. Lee Moffitt Cancer Center & Research Institute, an NCI designated Comprehensive Cancer Center (P30-CA076292).

## Supplementary Figures

**Figure S1.**
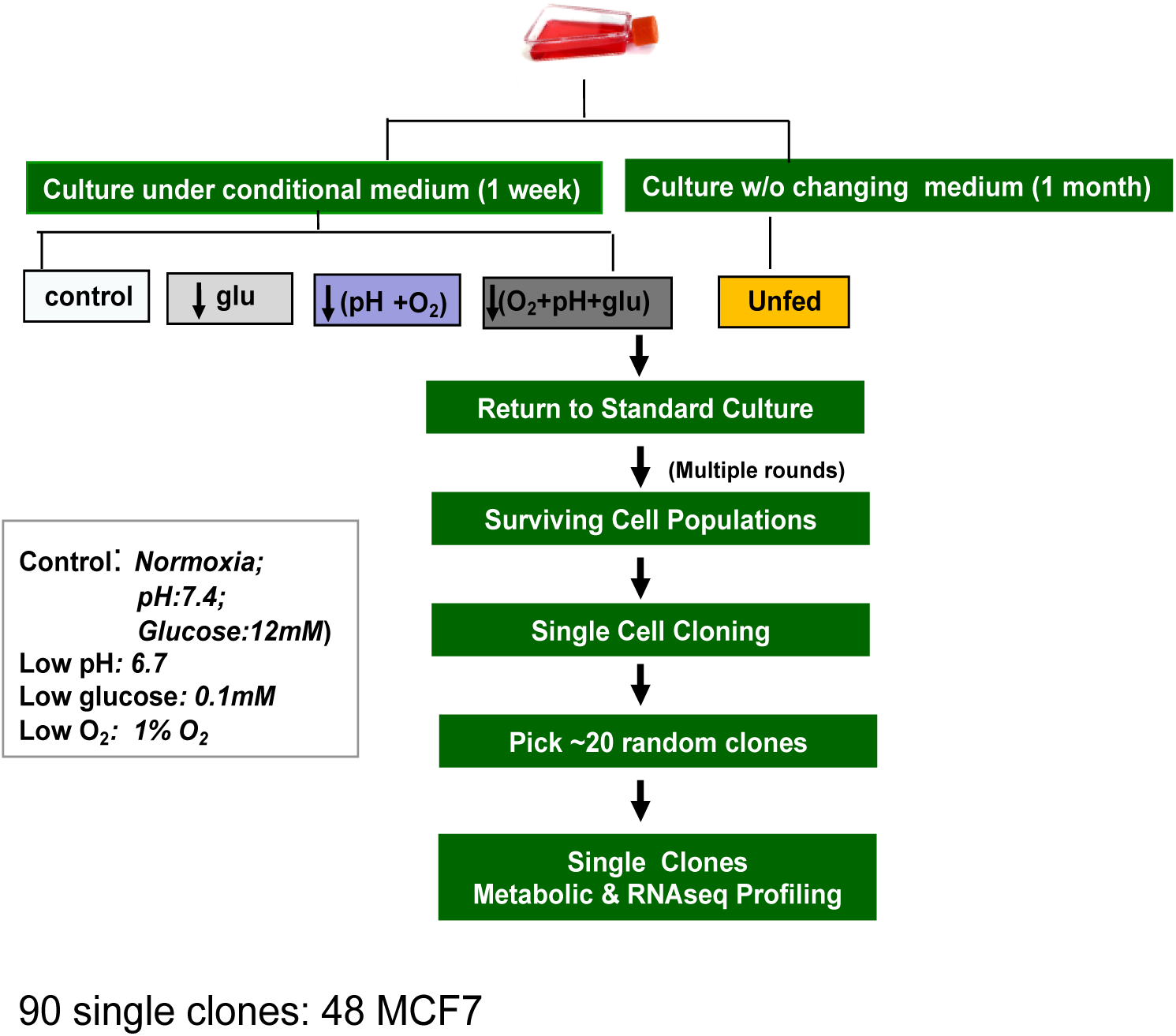
Experimental Design.

**Figure S2:**
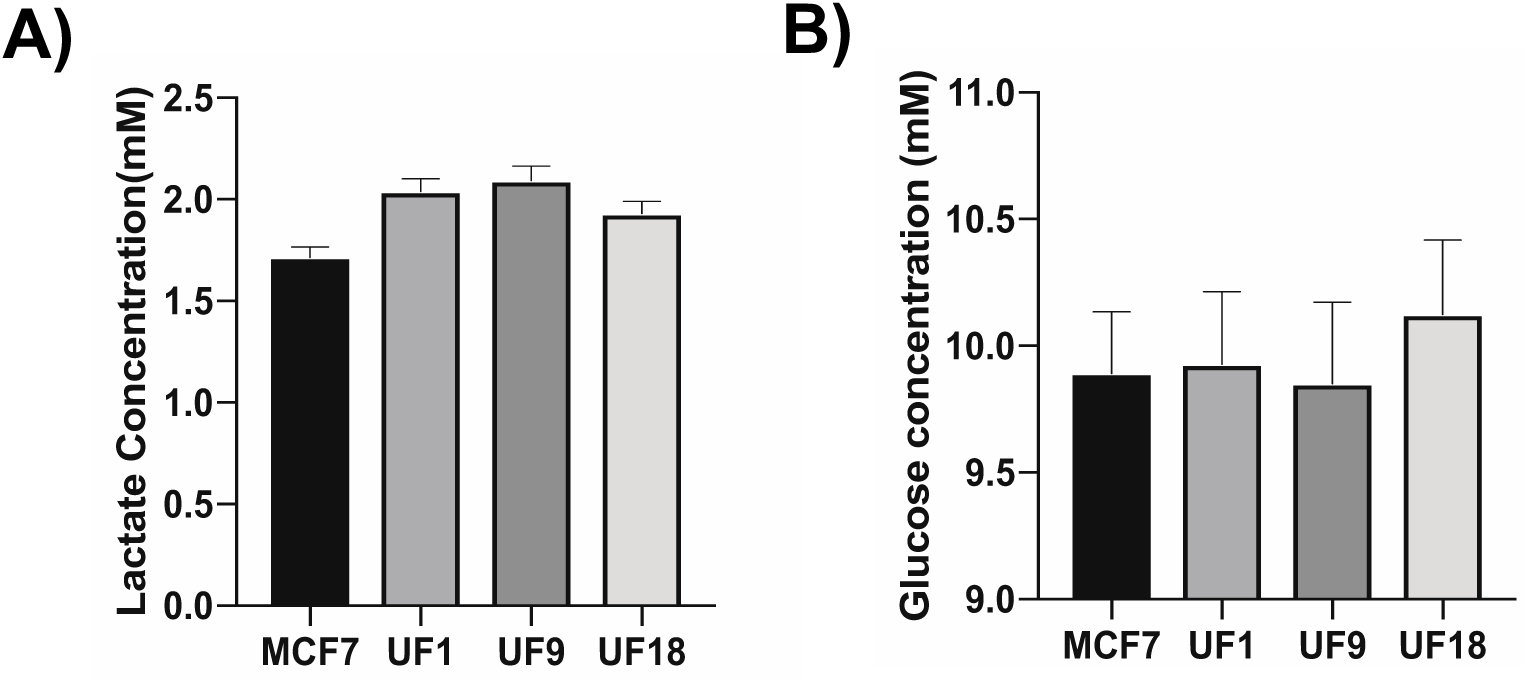
Unfed clones have Warburg phenotype. **A)** YSI measurements of three unfed clones and MCF7 cells. Unfed selected cells have higher lactate production rate.

**Figure S3:**
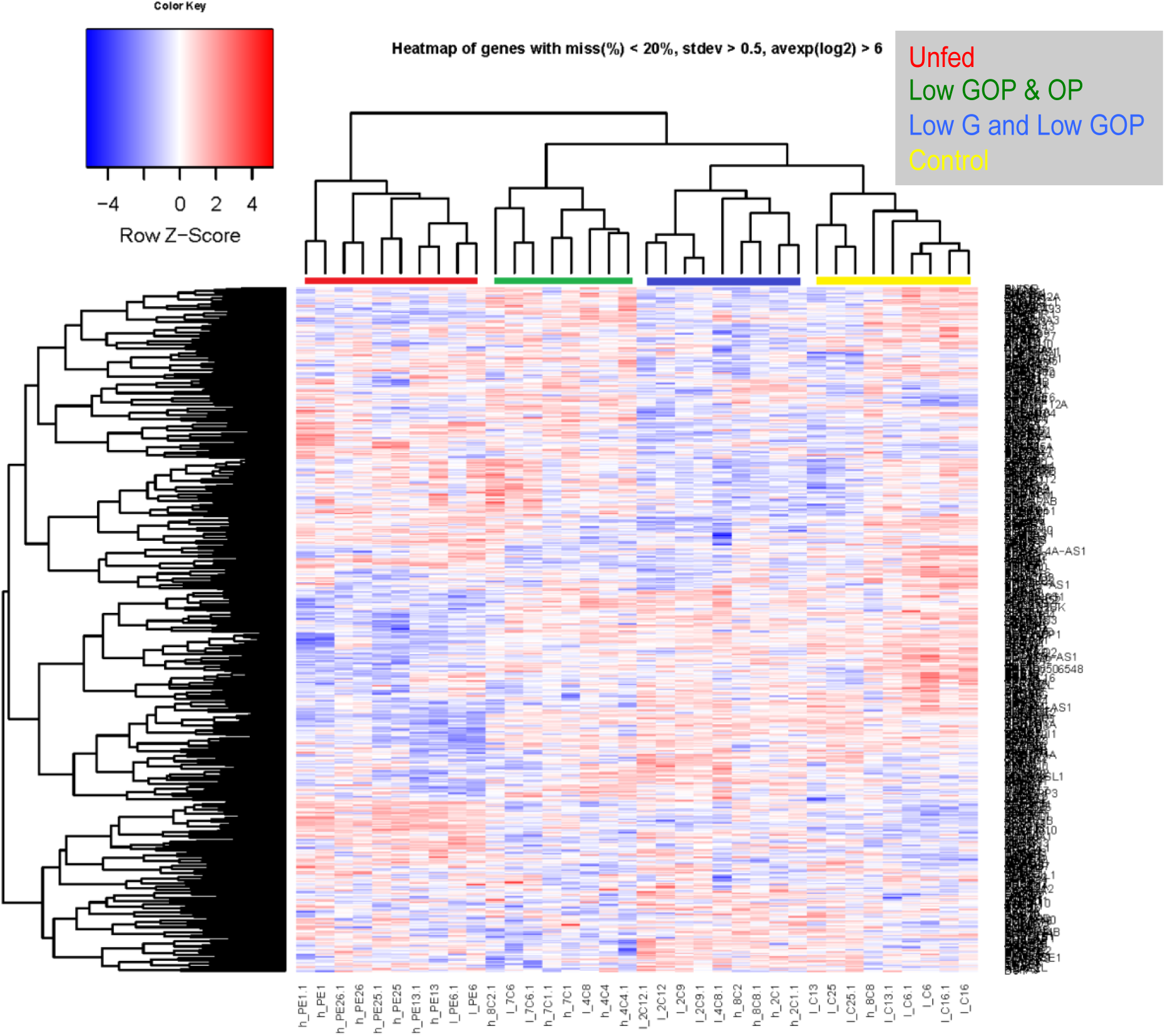
Different clustering RNA expression compared to Figure 2A. Within this one, the RED group consisted entirely of unfed cells; the GREEN group was comprised of GOP and OP cells, the BLUE group consisted of G and GOP cells, and the YELLOW group was almost entirely consisted of control cells. The clustering of cells from different selection patterns is not unexpected, as there is also great heterogeneity in the basal LPR between clones arising from the same selection conditions (cf. Figure 2). A preliminary analysis of RNASeq data was performed by linearly regressing gene expression data with lactate production rate and filtering for strongly correlated or anti-correlated genes. Unsupervised clustering (**Figure S3**) of these data showed that 1000 most highly correlated and anti-correlated genes clustered within selection condition. The red and green groups were the most glycolytic and selected by non-feeding and low Glucose, low Oxygen, and acidic pH, as the blue and yellow group were generally less glycolytic and selected with low glucose and low pH and oxygen.

**Figure S4.**
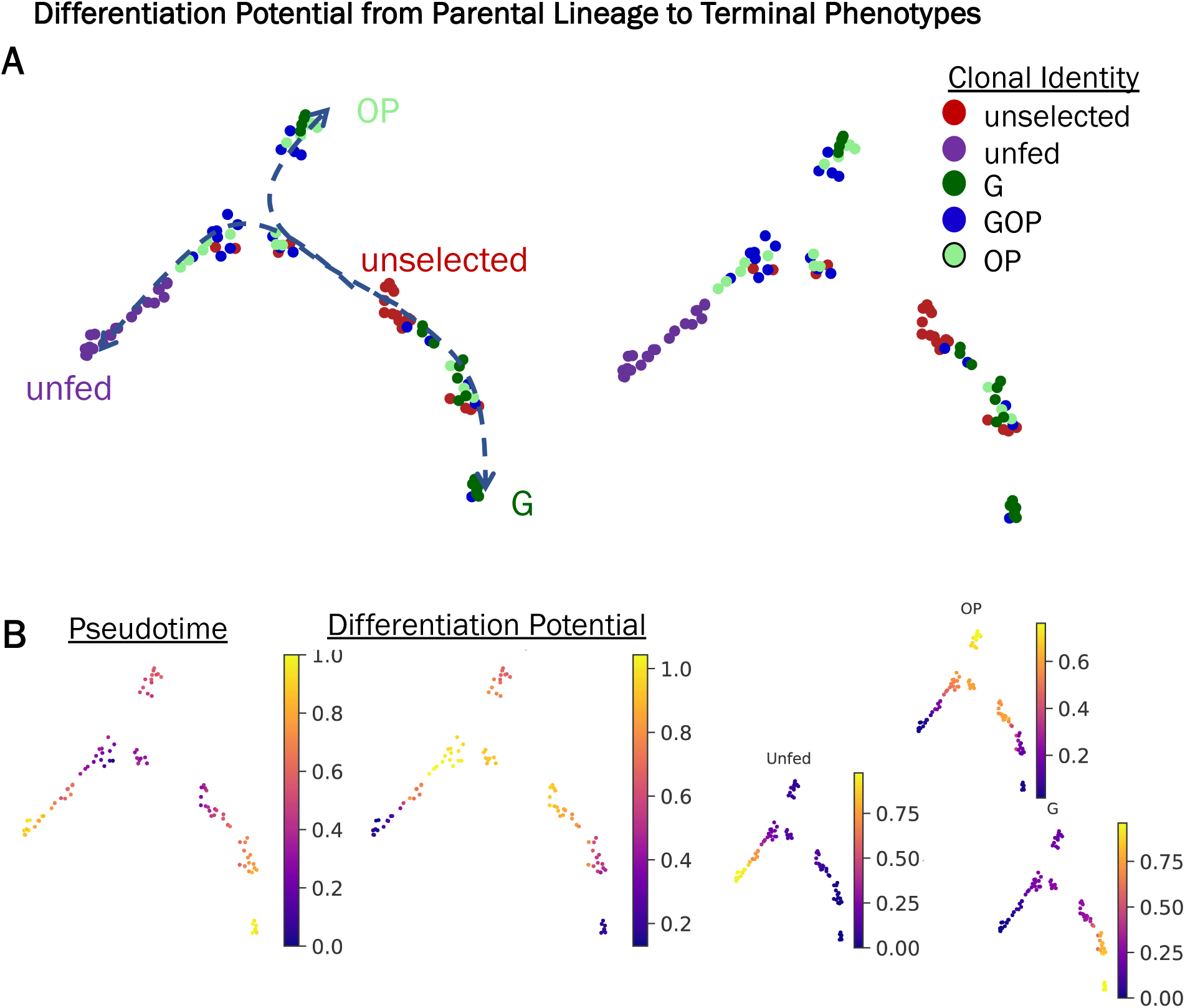
Phenotype evolution trajectory alignment of single clone RNA-sequencing for evolving breast cancer cell populations. (A) Cell fate analysis with Palantir was applied to the single clone RNA-sequencing dataset to determine differentiation potential from an initial, unselected, parental lineage to selected, phenotypic terminal states of G, OP, and unfed. UMAP projections were used to visualize the high-dimensional dataset and known identity of each clones was colored on the UMAP projections. Unselected clones were indicated in red, unfed clones were indicated in purple, G clones were indicated in green, OP clones were indicated in mint, and GOP clones were indicated in blue. (B) Pseudotime alignment and differentiation potential for each clone was calculated and colored onto the UMAP projection. Pseudotime ranged from 0, indicating earlier timepoints in the lineage trajectory in dark blue, to 1, indicating later timepoints in the lineage trajectory in yellow. Differentiation potential indicated the probability that a given clone would proceed along the trajectory. Differentiation potentials near 1, indicated in yellow in the plot, represented clones with the highest potential to proceed to a terminal phenotypic state. Differentiation potentials near 0, indicated in dark blue, represented clones already in a terminal phenotypic state and thus they had low potential of changing from that state.

**Figure S5.**
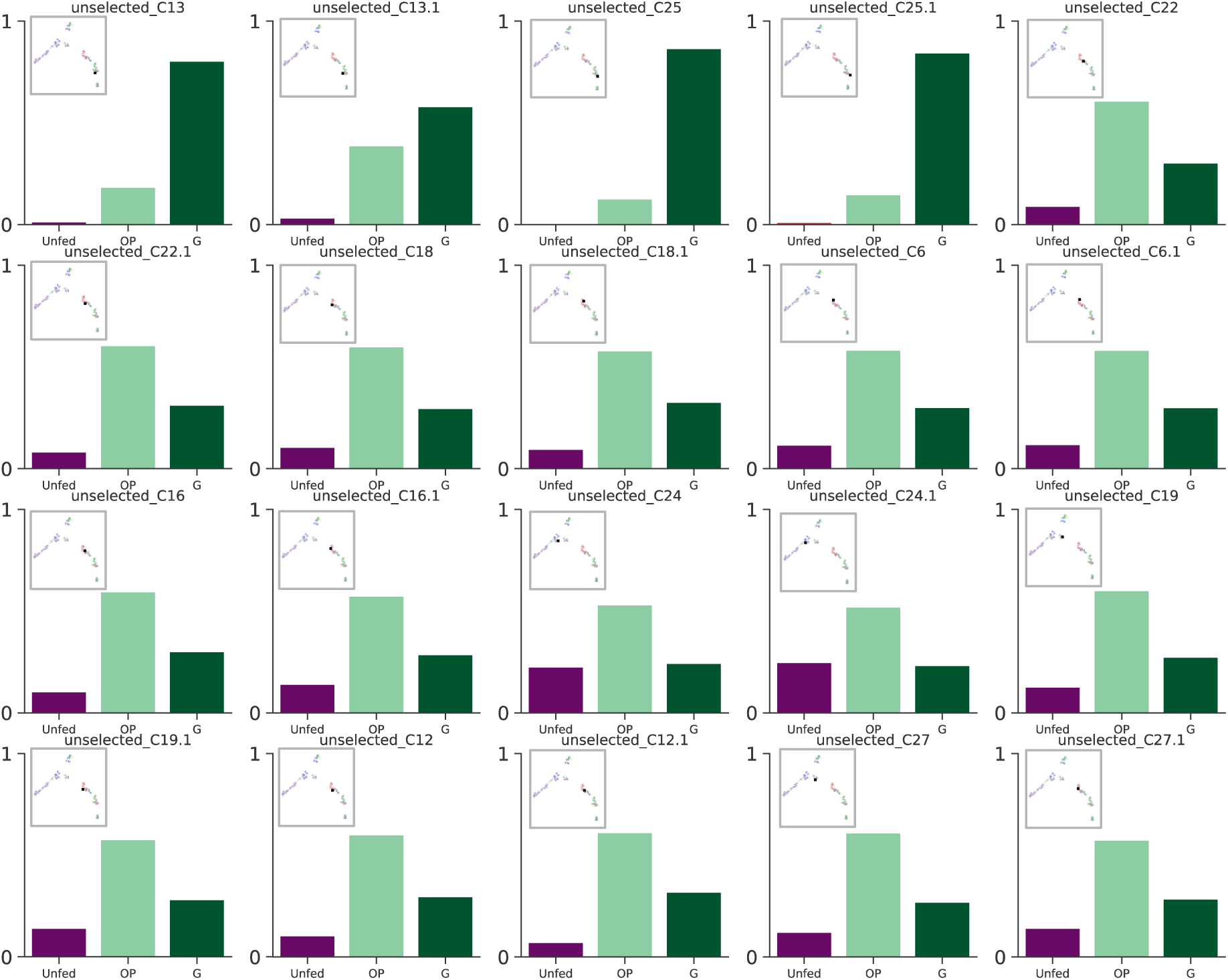
Terminal state probability distribution of individual unselected, parental lineage clones to terminal phenotypes: unfed, G, OP. For each unselected clone, the probability of that clone of evolving to a G, OP, or unfed phenotypic state. Four of the clones have increased probability of becoming G, while remaining sixteen have highest probability of becoming OP overtime. Interestingly, clones most likely to become G also have almost no chance of becoming unfed phenotype. Inlet shows the location of the given clone on the UMAP projection in Figure 2C and Figure S4 A and B.

**Figure S6:**
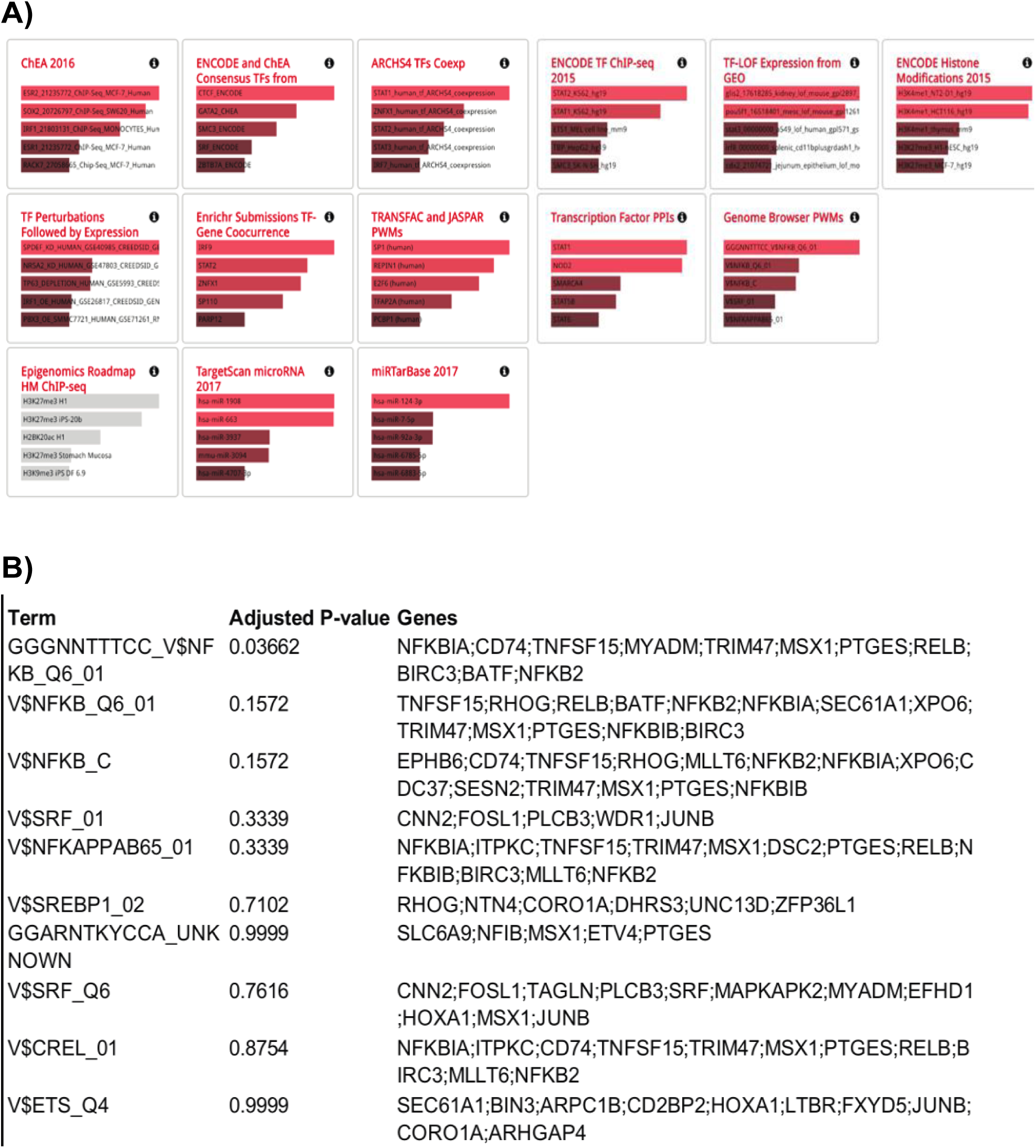
A) Transcription factor analysis with Enrichr (positive coefficient genes). We Used “Enrichr” (http://amp.pharm.mssm.edu/Enrichr/) for transcription factor enrichment of significant MCF7 genes B) Transcription factor analysis with Enrichr (positive coefficient genes).

**Figure S7.**
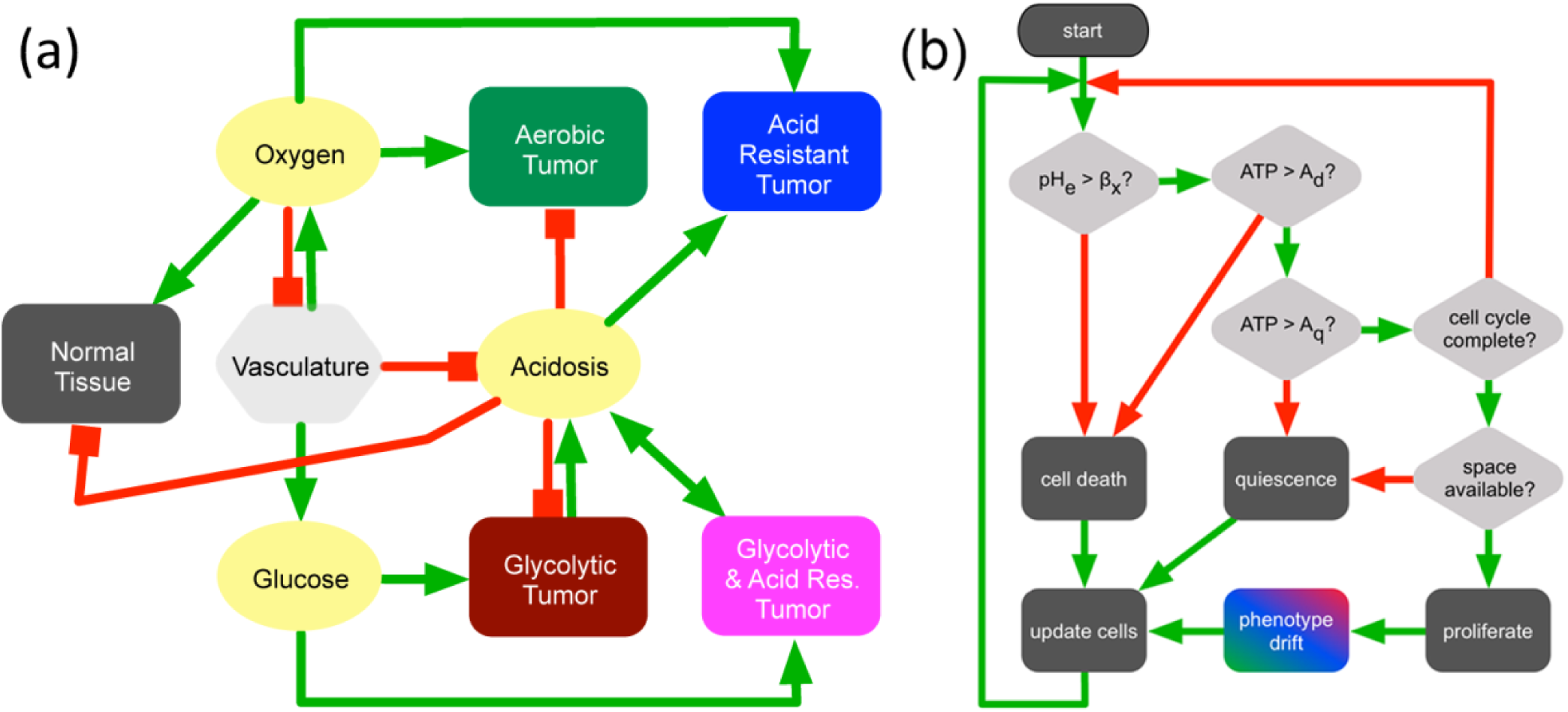
Schematic of the hybrid mathematical model. (a) The interactions between cells, vasculature, and metabolites in the *in vivo* simulations. Green lines indicate promotion or production; red lines indicate inhibition. Four generalized tumor cell phenotypes are shown, but the model has a continuous phenotypic variation. The pink cells are representative of the WE phenotype. (b) decision path for the model. Green lines are ‘yes’, red lines are ‘no’. Cells die if they have low ATP efficiency in the given conditions, or if they are maladapted for the level of acidosis. Cells change their phenotype by a small, random amount upon proliferation.

**Figure S8.**
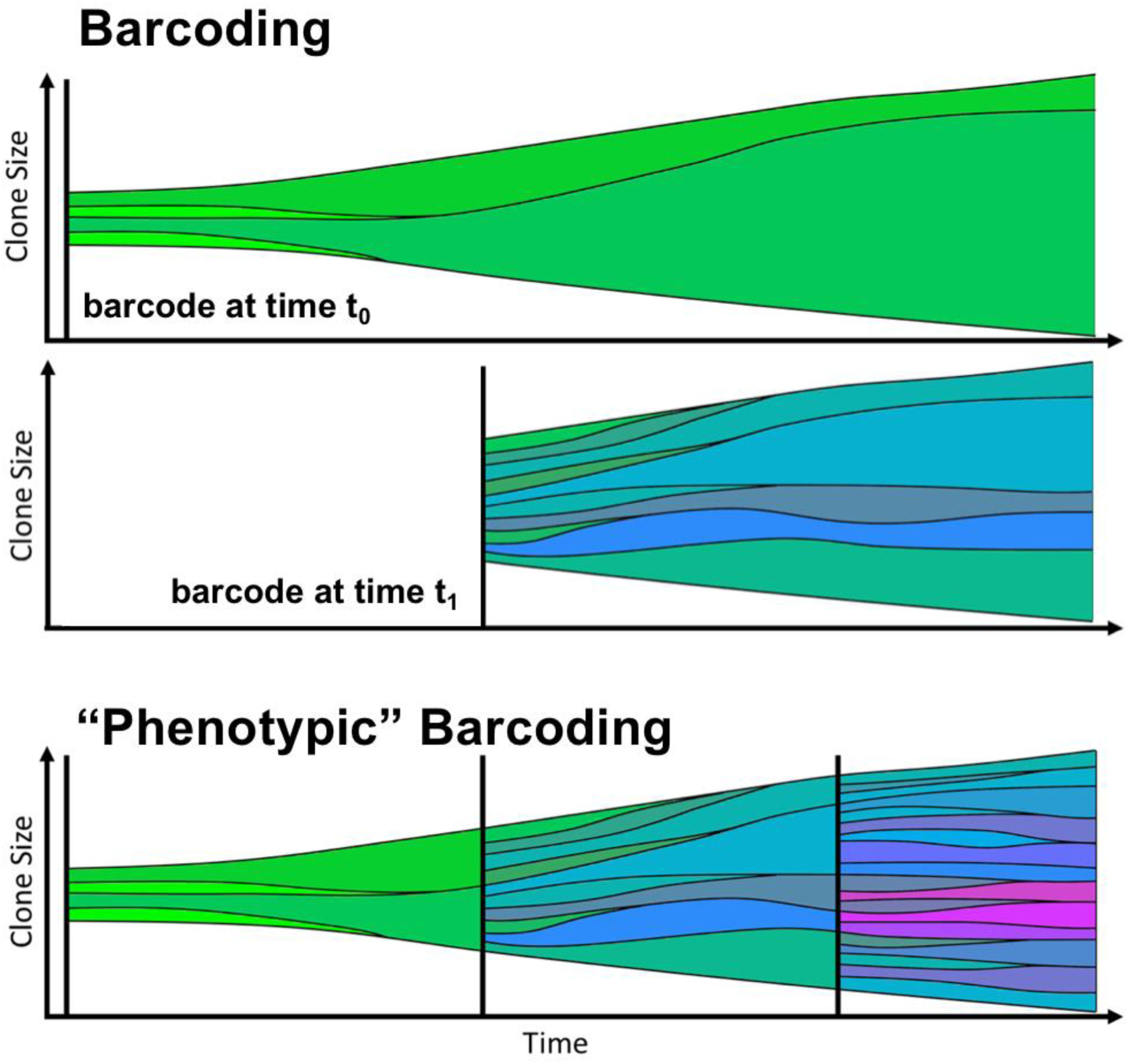
Illustration of barcoding scheme for the mathematical modeling. At time 0, all cells are given a unique ID (top). This can be repeated at later times (e.g., t_1_) by adding a second unique ID to each extant cell (middle). The clones and subclones can then be colored by average phenotype (bottom) so that the lineage of the final phenotypes can be determined.

**Figure S9.**
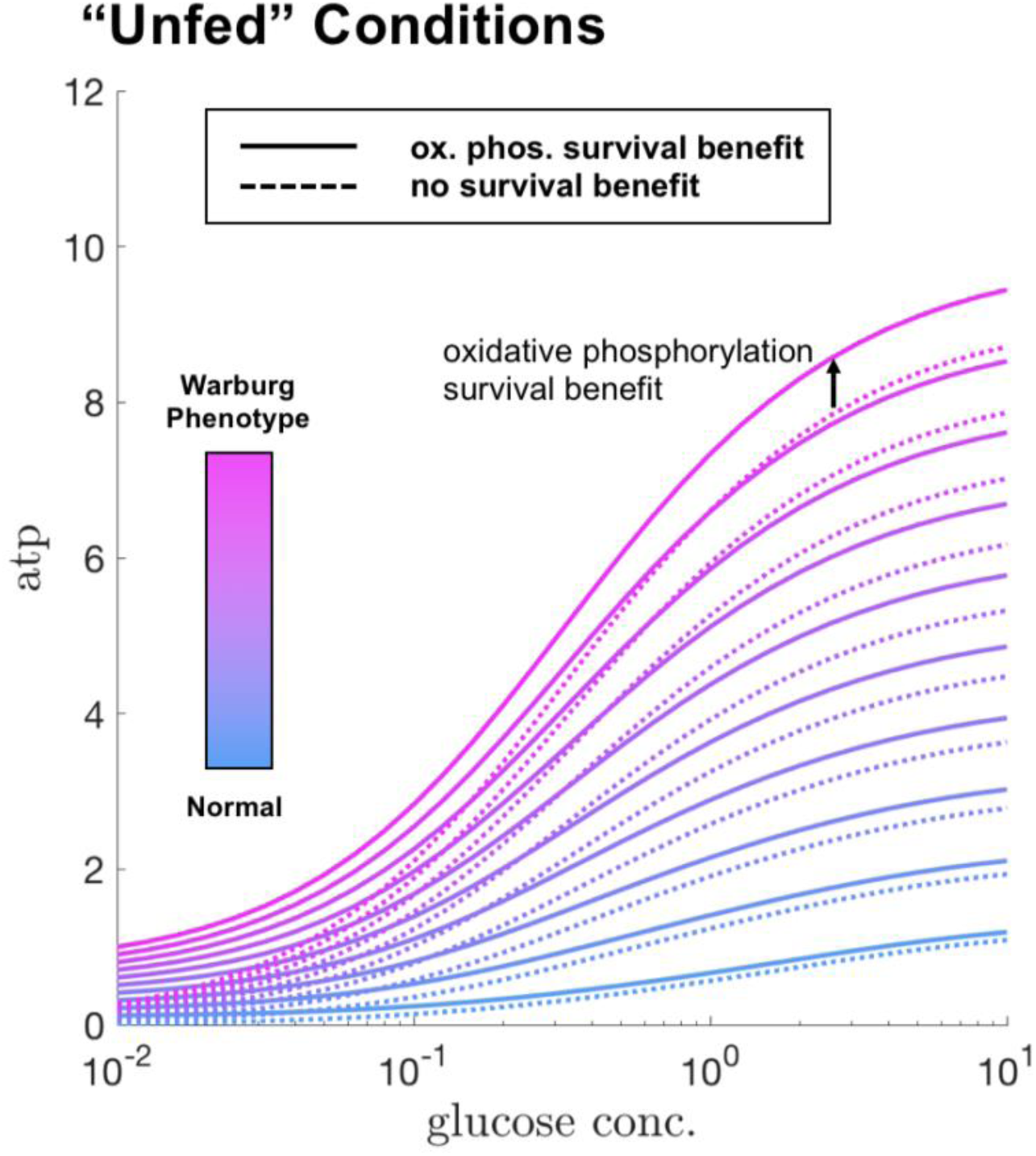
Quantification of the ATP efficiency benefit gained by the WE phenotype under different conditions of glucose, from the mathematical model results. The dashed lines show the original model ATP efficiency (vertical axis) for different concentrations of glucose. The solid lines show the efficiency with the addition of the survival benefit term. In all cases, this enhances the cellular ATP production, but WE phenotype cells gain the most benefit, particularly in poor conditions.

**Figure S10:**
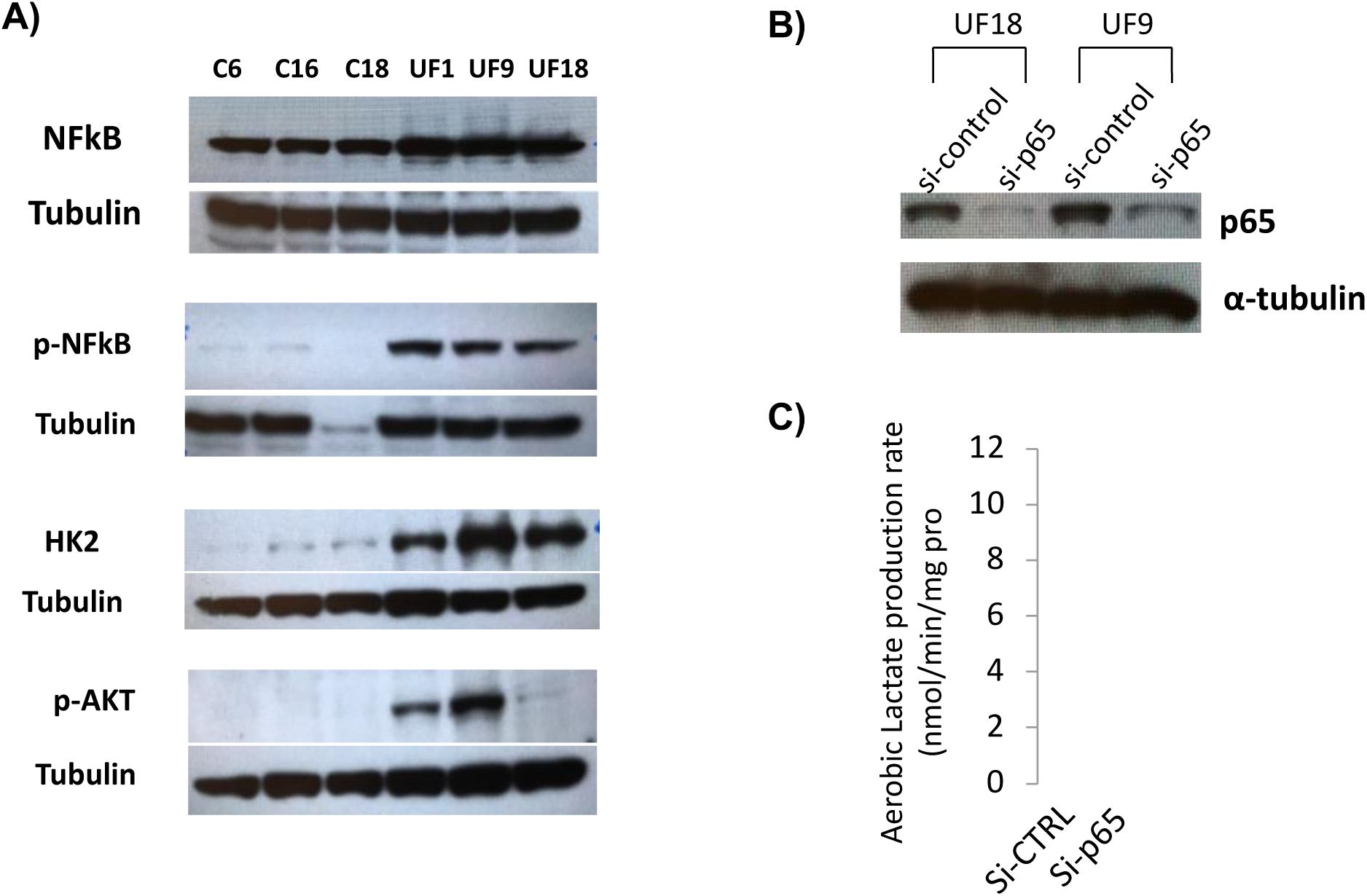
A) Western blot validation of NFкB and its clients. Unfed clones have higher NFкB, p-NFкB, HK2, p-AKT expression compared to the parental MCF7 clones. **B and C)** Knock down of NFкB reduces the glycolytic phenotype of unfed clones. B shows the validation of NFкB siRNA in reducing the protein expression. Using validated NFкB siRNA from B, we showed that lactate production rate of unfed cells decreases in C. Data are presented as mean with standard deviation as error bars.

**Figure S11:**
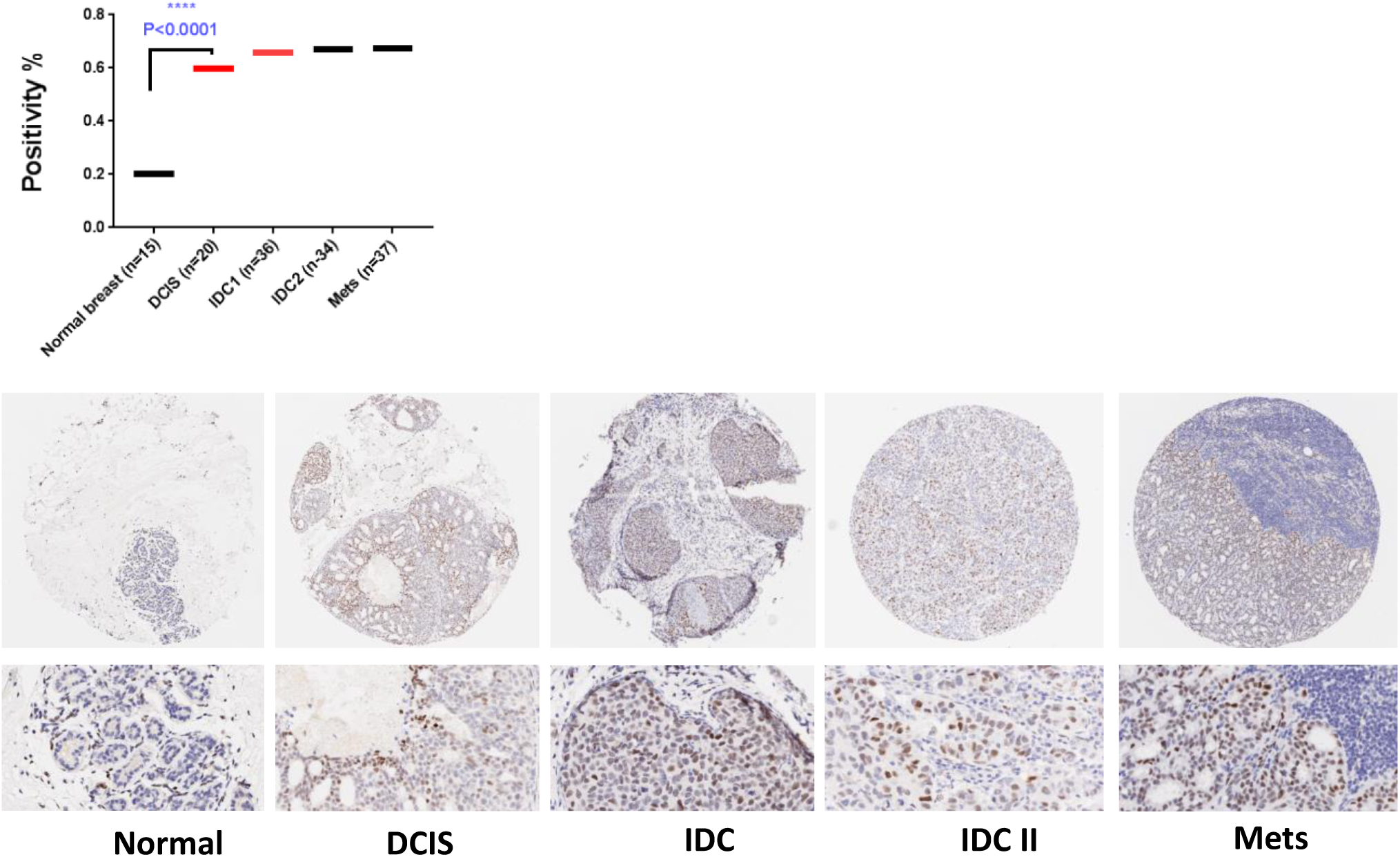

## Notes

### Summary of Updates

There was typo in the title that is fixed now!

